# Slow and *ad hoc*: Unravelling the two features characterizing the development of bacterial resistance to membrane active peptides

**DOI:** 10.1101/614370

**Authors:** Ayan Majumder, Meher K. Prakash

## Abstract

Membrane disrupting drugs such as antimicrobial peptides are being considered as a solution to counter the problem of antibiotic resistance. Although it can be intuitively imagined that bacteria will eventually develop resistance to this class of drugs as well, the concern has largely been ignored. Drawing upon the experimental data from the resistance of *Staphylococcus aureus* to antimicrobial peptides, we theoretically model the membrane adaptation under drug pressure. Using our model, we simulate the serial passage experiments with and without the drug pressure, and use the comparisons with experiments to estimate the unknown kinetic parameters. While the development of resistance to enzyme or membrane targeting drugs are both driven by spontaneous mutations, an additional lysylation step required in the latter slows the development of resistance. By quantifying the tradeoff between the gain in fitness under drug pressure and a loss in growth due to membrane modification, our model shows a fast reversal of membrane composition in drug free conditions, re-sensitizing the bacterium to the drugs.

## 1 Introduction

The discovery of penicillin dramatically increased survival rate by lowering mortality. Almost hundred years after this discovery, the world health organization (WHO) claims that antibiotic resistance is one of the major health threats of the 21st century. A rapid increase of bacterial resistance against almost all clinically used antibiotics is creating a real threat towards human health.^1^ For example, the treatment of *Staphylococcus aureus (S. aureus)* infections has become problematic due to the evolution of multi drug resistant strains. Daptomycin, a cyclic lipopeptide, which interacts with the membrane in a *Ca*^2+^ dependent mechanism was considered to be effective against *S. aureus*,^2^ but recent reports suggest the rise of daptomycin resistant strains. Similarly *A. baumanii* has also evolved into a multidrug resistant (MDR)^3–5^ bacteria which has already developed resistance against carbapenem,^6^ one of the last line drugs. Bacterial adaptation, primarily driven by mutations in the enzymes that are targeted by drugs, is the principal reason for this resistance. This crisis suggests a need to search for new antimicrobial agents, with non-conventional mechanism of action.

Antimicrobial peptides (AMPs) are found in almost all organisms^7^ and their widespread role in innate defence has been recognized.^8,9^ Antibiotics such as AMPs and their mimics which target bacterial membranes instead of their enzymes are being considered as effective alternatives to combat antibiotic resistance and many of them are already in clinical trials.^10^ Some AMPs target bacterial macromolecules^11^ like DNA,^12^ RNA^13^ or proteins.^14^ However, it is now accepted that membrane disruption such as by barrel stave^15^ and carpet^16^ mechanisms, is a key factor for AMP activity.^14,17,18^ Cationic AMPs are believed to be drawn to the membrane, mostly consisting of negatively charged phosphatidylglycerol (PG) and cardiolipin (CL), by electrostatic interactions, whereupon, hydrophobic forces play a key role in pore formation and membrane disruption.^19,20^ Since mammalian cells mostly consist of zwitterionic phospholipids, it is believed that AMPs do not act on them.^19,20^

Clinical introduction of membrane active drugs, including AMPs, is new. While it is commonly cited that bacteria do not develop resistance against AMPs,^21–23^ it is not technically true. Serial passage studies in which bacterial colonies are subjected to systematically increasing drug challenge, show a development of resistance, although it is slower than the conventional drugs. Several studies have shown that the bacterial membranes adapt, mainly by surface charge modification in the presence of AMPs. For example, *Staphylococcus aureus*,^24^ *Listeria monocytogenes*,^25^ *Mycobacterium tuberculosis*,^26^ *Bacillus anthracis*^27^ and *Rhizobium tropici*^28^ have enzymatic mechanisms to add positively charged lysines to the phospholipids.

Specifically in *Staphylococcus aureus* (*S. aureus*), the antimicrobial peptide-sensing (aps) system in the membrane senses the presence of AMP, which activates the Multiple peptide resistance factor (MprF). MprF catalytically mediates the reaction of converting a fraction of PG to lysyl-PG (LPG), and thus increases the positive charge of the membrane and repels AMPs, as shown, for example, in the development of resistance to daptomycin.^29,30^ Mutant *S. aureus* with out the capability of LPG production shows reduced virulence in *in vivo* studies.^31^ The development especially of MRSA in *S. aureus* has been noted with the upregulation of MprF^32–34^ and/or the single nucleotide polymorphism (SNP) in the MprF open reading frame (ORF)^32,35,36^ and a consequent increase in the LPG component of the membrane. A single mutation in the MprF, such as the most common L826F^37^ is sufficient to activate the enzymatic conversion of PG to LPG, and hence can contribute towards the resistance against AMPs. However, as may be expected, the kinetics details of the mechanisms are not completely known.

It thus becomes relevant to characterize the nature of the drug resistance that evolves in this new strategy and to quantify the rates of development and reversal of resistance to AMPs. While theoretical models have probed the rates of evolution of resistance against conventional drugs,^38^ thus far, to our knowledge there is no quantitative model that delineates the evolution of resistance in a bacterial population in the presence of AMP. In this work, we develop a computational model for the development of resistance by surface adaptation. Possible resistance to peptides by alternative mechanisms such as the action of peptidases, are usually considered secondary, and are not considered in this work. We model the stochastic appearance of mutations that will upregulate MprF and generate LPG by catalytic lysylation, in addition to the basal rate of LPG production. The majority of assumptions, including the membrane composition and the increase of MIC indicating the development of resistance in serial passage tests^29^ are drawn from the data on CB118 strain of *S.aureus*, as this was the most comprehensive data we could find. However, the conclusions are expected to be of general relevance to all membrane active drugs.

## 2. Model assumptions

### 2.1 Growth in the absence of drugs

The CB118 strain of *S. aureus* membrane is mainly composed of PG, LPG and CL, in a proportion of 83.96%, 12.37% and 5.38% respectively.^29^ *S. aureus* was assumed to be spherical with a 1 *μm* diameter and that the area of each molecule of PG, LPG and CL is 60, 60 and 120 Å^2^ respectively. We estimated the number of PG, LPG and CL in a single cell of *S. aureus* to be 7.8 × 10^6^, 1.2 × 10^6^ and 5 × 10^5^ respectively. In a growth medium and in the absence of antimicrobial drugs, the total phospholipid increases at a constant rate, following zeroth order kinetics, while the composition of the different lipids remains almost constant.^39^ In our model, a bacterial cell is assumed to divide when the number of phospholipid molecules double. Using these parameters and Gillespie algorithm,^40^ we modelled the growth in a passage and reproduce a typical experimentally observed growth curve. The rate of formation of lipids obtained from Short *et. al*.^39^ was corrected by a factor of 0.55 to obtain a cell-division rate of 25 minutes^41^ in the exponential growth phase 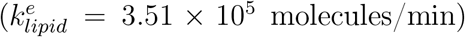. The rates of formation of PG 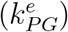, LPG 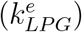 and CL 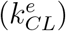 in the exponential growth phase were obtained by multiplying 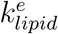 with the corresponding percentage composition, *i.e*., 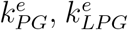 and 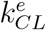 are 2.95 × 10^5^, 0.43 × 10^5^ and 0.19 × 10^5^ molecules/min respectively. The growth phase dependent rate of bacterial lipid synthesis at a time *t* (in minutes) during a serial passage is obtained by multiplying the rates in the exponential phase with a phase dependent gaussian factor (**Supplementary Figure 1**): *f_PhaseFactor_* = (0.07 + *exp*{6 − (*t*/120)}^2^). The choice of the gaussian for the rate comes from the derivative of sigmoidal growth of bacterial numbers, and its width was chosen to match a factor of 10^4^ gain in colony size during the 24 hour serial passage experiment.

### 2.2 Effect of drug on bacterial death and growth

#### Increase in LPG production

In the presence of AMPs, in addition to the biosynthesis of PG, CL, LPG, an additional enzymatic conversion of PG to LPG is activated. We assume that an SNP in MprF that is required to drive this conversion happens in 1 out of 10^7^ cell divisions and continues to be present under drug pressure. The rate of PG conversion gets upregulated with the increasing MprF levels. We assume an upregulation factor of *f_MprF_* after each cell division since the mutation, increasing by a factor (*f_MprF_*)^*N*^ over *N* cell divisions 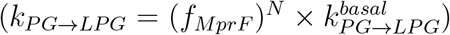. The only *k_PG→LPG_* that was measured^42^ was for *E. coli* (0.055min^−1^) rather than *S. aureus* and after the bacteria adapted to a high concentration of the drug. We optimize the 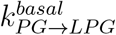 to be used in our model, as well as the multiplicative factor *f_MprF_, a posteriori* by comparing the results from our model with the experimental data (**Results Section**).

#### Fitness cost

Gaining antibiotic resistance by gene mutation is associated with a fitness cost, *i.e*., the resistant bacteria has lower growth rate than the susceptible strain.^43–47^ Susceptible strains maintain homeostasis in the cell, aided by the membrane phospholipids. Under drug pressure, when the membrane composition changes, it affects the growth rates. When bacterial enzymes are targeted, the fitness may be regained over the generations, with compensatory mutations. However, the same is not true when the membrane undergoes a surface charge modification. We assume a linear drop in fitness, *f_fitness_* = 1 − {(*f^max^*/12) × (%*LPG_Mutant_* − 12)} (**Supplementary Figure 2**), where *f^max^* is the maximum drop in fitness when the lysylation is at the maximum level (24% LPG in the membrane) as observed in the *S. aureus* data and the value of *f^max^* that was determined by comparing the results from model with the experimental data (**Results Section**). This correction factor was applied to the growth rate at different membrane compositions. However, we found with our calculations that the general conclusions were not sensitive to small changes in the estimates of the two factors, *f^max^* and *f_MprF_*.

#### Death rate

Unlike enzyme active drugs which are inactive in the stationary phase, AMPs continue to act depending only on the membrane composition rather than the growth phase. Motivated by the data from the CB118 strain of *S. aureus*, we assume the death rate is highest at 12% LPG and drops to zero beyond 24% LPG. In our model, we use a sigmoidal dependence of death factor on the %LPG, *d_Sigmoidal_*(%*LPG*) = *exp*(−0.9 × (%*LPG* − 18)) /1 + *exp*(−0.9 × (%*LPG* − 18)) (**Supplementary Figure 3**) to model this gradual loss of sensitivity to drug as the membrane composition changes.

## 3 Results

### 3.1 Simulating serial passage

Each passage in our simulation begins with a colony of 10^9^ bacterial cells and grows over a period of 24 hours, during which the colony grows to a size of 10^13^ (approximately). A schematic of how the bacterial resistance develops or is lost in our model is shown in **Figures 1a and 1b** respectively. It is assumed that mutations in MprF occur at a rate of 10^−7^ with every cell division, and the mutant MprF upregulates at a fixed rate under drug pressure, which in turn affects the LPG levels. A heterogeneous mix of sensitive and resistant strains developed under drug pressure is used to inoculate the next passage. In addition to the formation of the lipids, as noted in Section 2.2, the MprF dependent rate of modification of PG to LPG is also implemented in our model. We determine this MprF dependent rate *a posteriori* by comparing our results with the experimental data. Using phase and LPG dependent factors, the rate of increase in the number of lipid molecules is given by 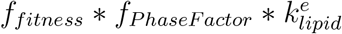, and the logarithm of the fraction of bacteria that die over time *t* (min) as {*d_Sigmoidal_*(%*LPG*) × {(*t*/(5 × 24 × 60)) × log(10^10^)}. Bacterial variants at any stage during cell divisions, can be described as *B*(*LPG, MprF*) by the levels of LPG and MprF. %LPG defines the resistance and death rates, and the MprF indicates the rate of modification of PG to LPG. MprF and LPG thus act analogous to descriptive parameters in mechanics - velocity and position. Similarly when the AMP pressure is relaxed, bacterial composition slowly reverts back to the sensitive one. As different types of mutant cells with varying %LPG composition produced after each susceptible cell division, we bin the mutant strains in increments of 2% LPG, as R1, R2, R3…,R7 from 12% to 26%. MprF corresponding to the arithmetic average of the number of PG, LPG, CL of all the strain in a bin was used as the MprF in the bin. At the end of a passage, from the approximately 10^13^ bacteria, a sample of 10^9^ proportionate across the resistant strains is proportionately and used as a seed for the next passage.

**Figure 1:**
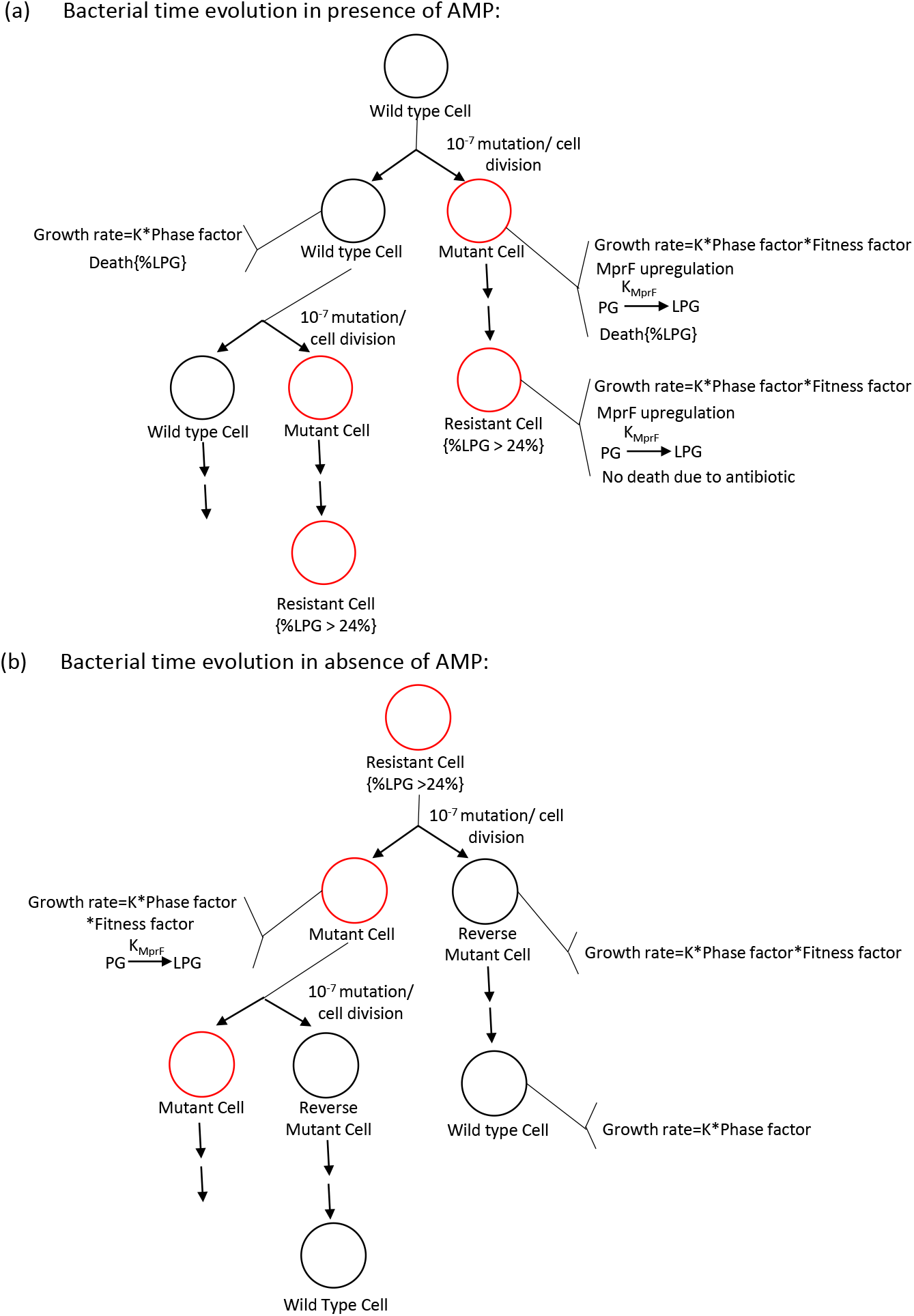
Schematic of the Model used in our simulations. (a) Mutations develop randomly, and lead to a conversion of PG to LPG. In the presence of an AMP, these mutants with modified surface charge have a fitness advantage. (B) In the absence of AMP, homeostasis requires the reversal of the membrane lysylation. A drop in MprF along with the mutations causes the LPG percentage to be reduced to the level of wild type. The black and red circles represents the presence of wild type MprF and SNP in MprF respectively.

### 3.2 Upregulation of MprF and evolution of resistance

MprF upregulation is very important for the adaptation of *S. aureus*. If *k_PG→LPG_* were to depend only upon the presence of MprF mutation, the LPG level in the mutants saturates in half of a passage time (**Supplementary Figure 4**), irrespective of how low the rate is. The experimental data on CB118 strain,^29^ suggests a longer saturation time, which can be achieved if MprF starts with a base level value 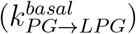 and upregulates after every cell division. In our model, we increment this rate by a factor *f_MprF_* which represents the increase of the MprF over every cell division. We parametrically scan through different MprF upregulation factors **Figure 2**. Comparing with the experimental data on the increase of LPG, we find *f_MprF_* = 1.009 after each cell division best models the cell division. The overall upregulation ((*f_MprF_*)^*N*^) after *N* cell divisions and %LPG with number of cell divisions are given in **Supplementary Figure 5**. The study of bacterial colony in the presence of AMP is given in **Figure 3.**

**Figure 2:**
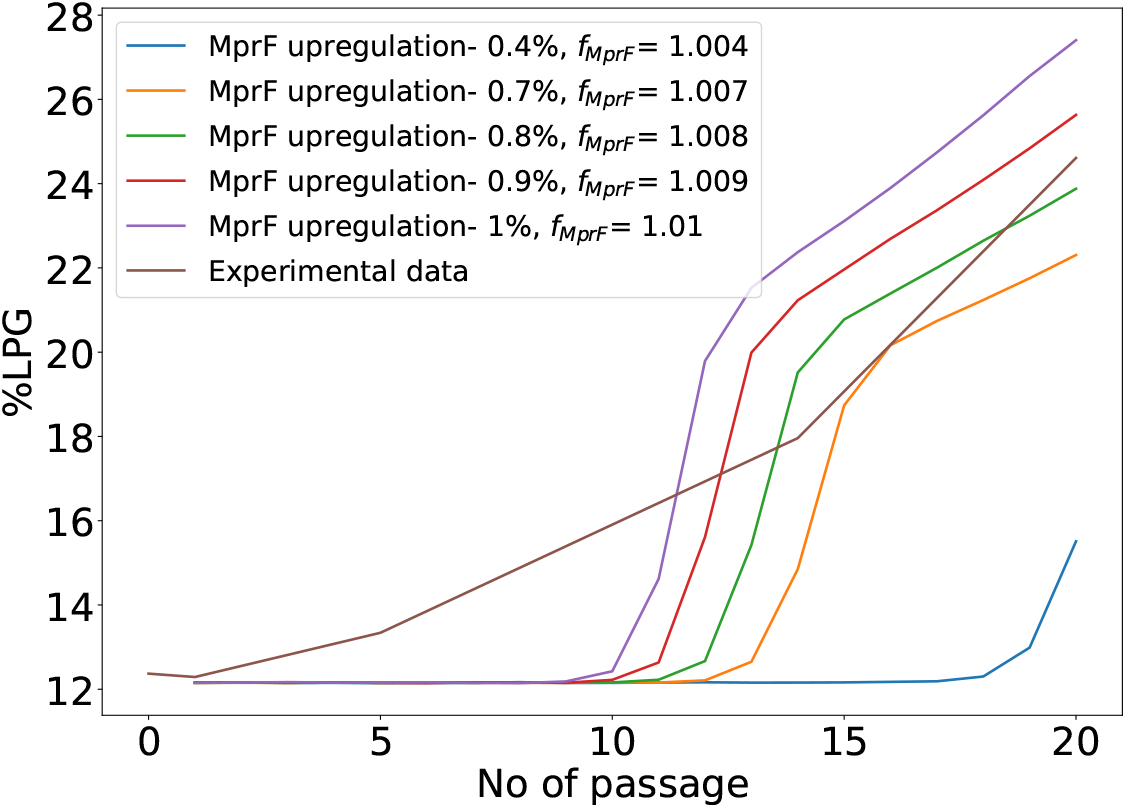
Change of %LPG of a mutant cell over a 20 day serial passage in presence of AMP, by considering the upregulation of MprF after each cell division. For this simulation the *f^max^* factor in the fitness considered to be 0.3, as justified later. Comparing the increase in LPG from theory and experiments, *f_MprF_* = 1.009 was used.

**Figure 3:**
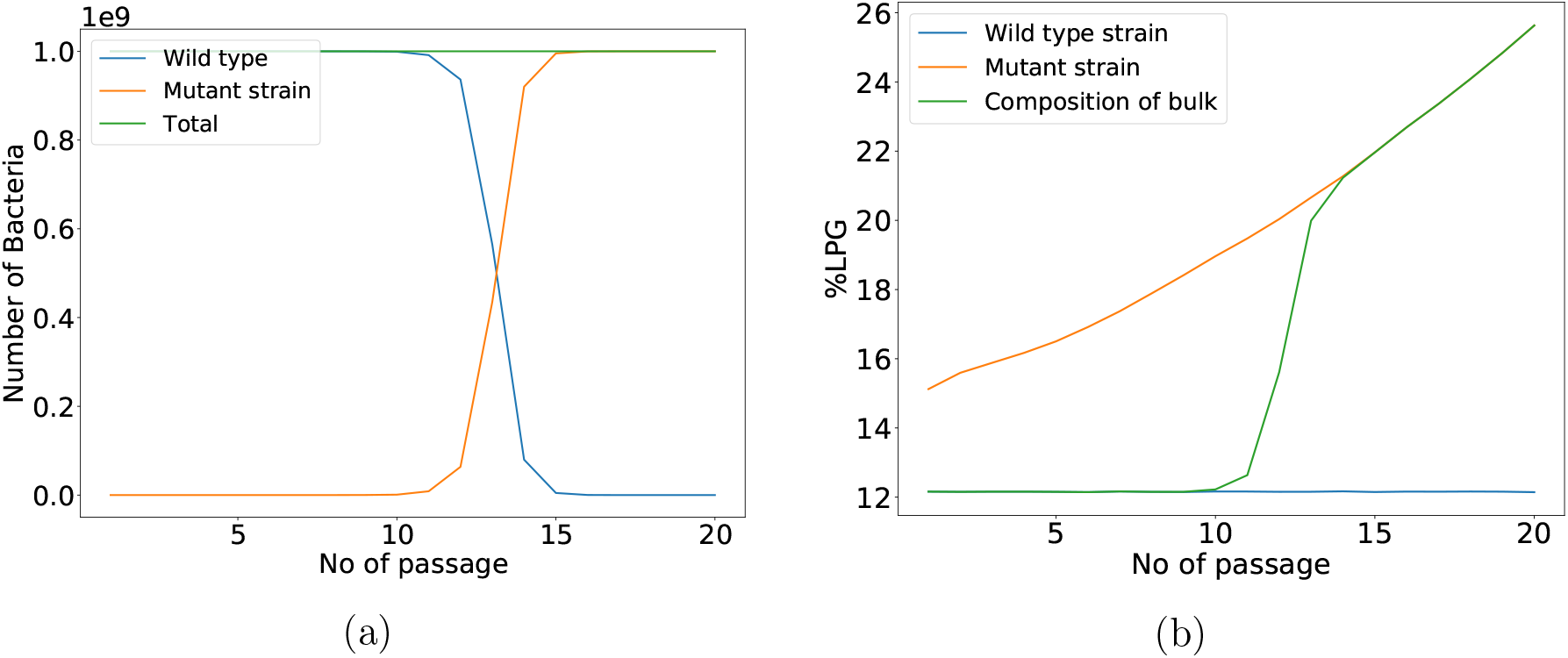
Time evolution of bacteria in presence of AMP, using *f^max^*=0.3 and *f_MprF_* = 1.009. (a) Change in the number of bacterial cells and (b) change of %LPG, over the passages are shown. Blue and orange lines represent susceptible and mutant cells, while the overall composition in the system are represented by the green lines.

### 3.3 Reversal of resistance upon AMP withdrawal

In order to understand how the bacteria behave when AMP is withdrawn from the medium, a colony starting with all mutant cells was studied for its evolution during the successive passages. Similar to the study under drug pressure, we simulated passages of 24 hours, with the last 5.5 hours being considered as a stationary phase. The two competing effects are the reversal of the mutations that were beneficial under AMP with a rate 10^−7^ and a reduced growth rate with the now higher LPG at no additional selective advantage for these species as the drugs are absent. After each cell division of the mutant cell, it produces a heterogenous distribution of the revertant mutant strains, which were binned in increments of 2% LPG as R1, R2,…, R7 for further analysis.

The revertant cells will have the higher fitness over the mutant cell. So, the revertant cells will divide faster than the mutant cells and eventually dominate the population. The number of days that the revertant cell will take to dominate over the mutant cell is depends on the fitness factor *f^max^*. We simulated 20 days of serial passage under the no-drug condition, and parametrically scan through different values of *f^max^* (**Supplementary Figure 6**). The results obtained as the change in the numbers of sensitive cells, and the overall %LPG in the medium are shown in **Figure 4**. This comparison was used to obtain *f^max^* = 0.3, which is used in the rest of our calculations.

**Figure 4:**
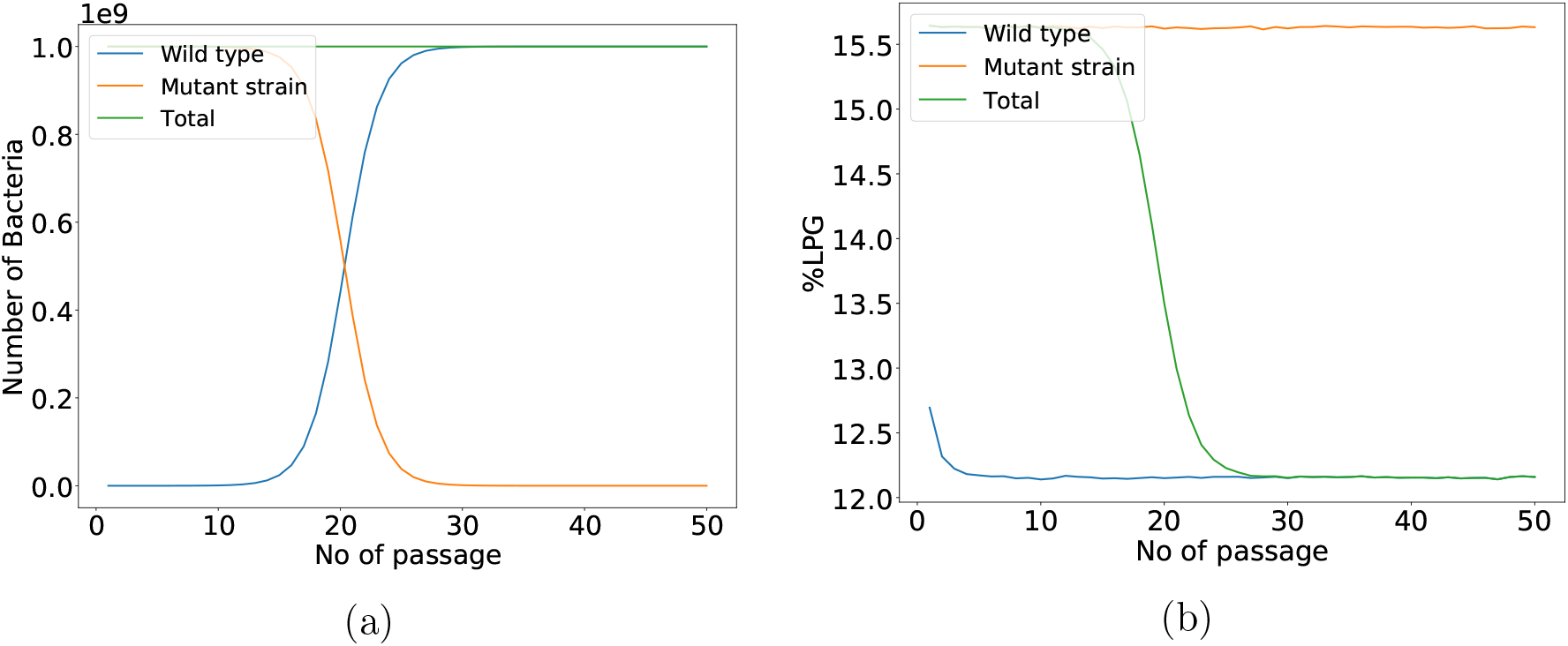
Time evolution of the bacteria in absence of AMP, using *f^max^*=0.3. (a) Change of number of the bacterial cells, and (b) change of %LPG over the passages are shown. Blue and orange lines represent susceptible and mutant cells, while the overall composition in the system are represented by the green lines.

### 3.4 Resistance to enzyme versus membrane targeting drugs

When a traditional drug hinders the activity of a bacterial enzyme, bacteria with mutations in the targeted enzyme get preferentially selected. Due to the mutations, a susceptible strain directly converts to the resistant strain and the fitness of the bacteria decreases rapidly. We simulated the difference in the patterns of resistance between enzyme and membrane targeted cases over time and the results are shown in **Figure 5**. A conclusion that emerges from this comparison is that the resistance to AMPs which is a two step process - an SNP is required to initiate the development of resistance, followed by an increase in LPG over time. The resistance to AMPs is thus not instantaneous even after the beneficial mutation in the bacteria. For an easy comparison, the gain of fitness in the resistant strain whether the enzyme or the membrane was targeted, was considered to be the same, *i.e*., the *f_fitness_* with 24% LPG under AMP drug pressure was used as the fitness of the resistant strain with mutant bacterial enzyme under the pressure of traditional drug.

**Figure 5:**
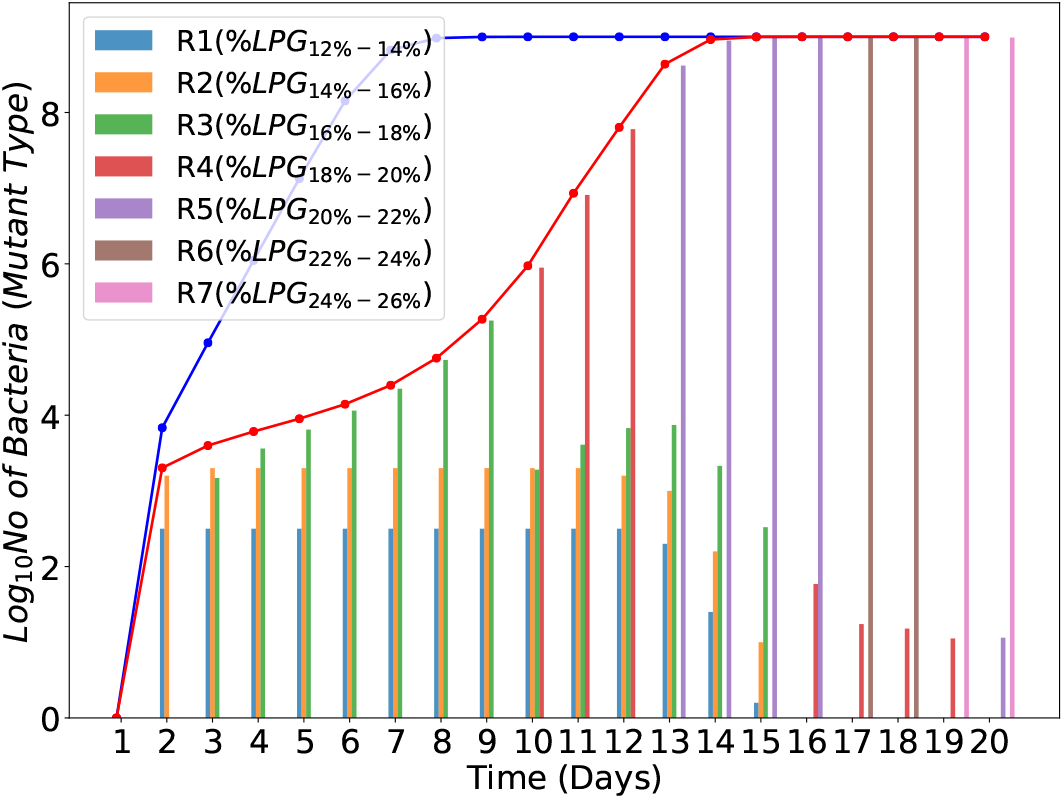
Comparison of the development of resistance in presence of AMP and traditional enzyme targeting drugs. The blue and the red curves represent the number of mutant cells in the presence of traditional drugs and AMP. The numbers from the heterogeneous mix of the strains R1 to R7, developing in the presence of AMPs is shown with the vertical bars. To simulate the time evolution of bacteria in presence of AMP, we considered *f^max^*=0.3 and *f_MprF_*=1.009 and to simulate the time evolution of bacteria in presence of traditional drug, we consider the fitness drop same as in case of AMP, *i.e*., 0.3.

### 3.5 Bacterial response to sudden removal and reintroduction of AMPs

Some studies previously reported that the antibiotic resistance remains persistent due to the compensatory mutation.^43,45,48^ But this is not the case for all antibiotics, as other studies established the reversion of resistance. Methicillin resistant strain of *S. aureus* (MRSA) has lower fitness than the methicillin susceptible strains, and a rapid decrease in MRSA when the use of methicillin is reduced has been noticed.^49^ Because, this resistance carries a fitness cost, a successive switch on and switch off of the antibiotic treatment can be a proper way to treat bacterial infection without causing resistance.^47,50–53^ It is hard to establish a correlation between the serial passage tests which are conducted on the scale of days with extremely high drug doses and the development of clinical resistance which happens over years, which is usually attributed to poor adherence. While a population based model for resistance may be explored as a separate work, we use our model, to study the consequences of a cyclic dosage, where the antibiotic treatment is repeatedly switched on and off. Starting with the 10^9^ bacteria, we studied the fate of the bacteria in a successive switch on and switch off dosage with the antibiotics. The results from an overall 60 day serial passage test with alternating periods with and without antibiotics for 10 days each are shown in **Figure 6**.

**Figure 6:**
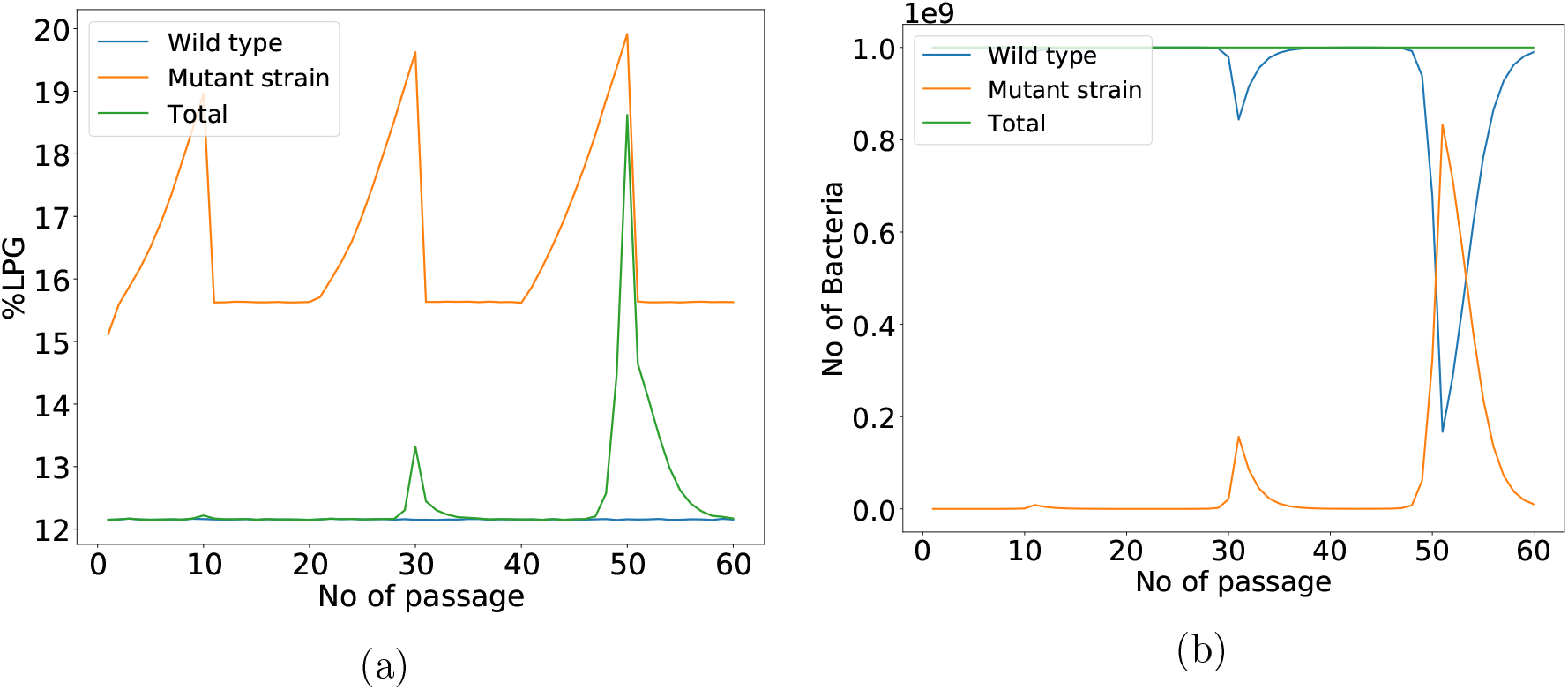
Fate of bacteria in a repeated switch-on and switch-off of AMP. (a) Change of %LPG, and (b) change of number of bacteria population over time are both shown. Blue and orange lines represent susceptible and mutant cells, while the overall composition in the system are represented by the green lines.

## 4. Discussion

### 4.1 Resistance is not instantaneous

When bacteria are challenged by a drug, targeting one of its critical enzymes, mutations which occurs rarely with a chance of 1 in 10^7^ offer drug resistance. But when they occur, the resistance to the bacterial cells is almost instantaneous, as the drug immediately loses its potency to bind the enzyme, thus losing its efficacy. The development of resistance against AMPs on the contrary happens in two stages. In the first stage, a mutation similar to the one in the enzymes targeted by the conventional drugs occurs. This mutation by itself does not provide drug resistance, but rather prepares it for resistance by charge modification. The next step is the upregulation of MprF which will increase the LPG levels, and thus over time resistance is achieved. This fundamental difference is responsible for the slower development of resistance in serial passage tests when using AMPs, typically requiring twice as many passages to reach the same level of resistance (**Figure 5**).

Another interesting aspect this two step process of drug resistance offers is that it raises the possibility of using a secondary drug target against MprF. For the next generation drugs, one can consider the MprF as a secondary drug target. MprF has two pockets binding PG and *tRN A^lys^* molecules respectively; and these could be targeted to inhibit lysylation.

### 4.2 Resistant strain is not unique

Since the development of resistance happens over several cell-divisions as an increment in the % of LPG, there are several intermediate strains with varying levels of MprF and resistance. This is unlike the case while targeting enzymes, when the resistance strain is unique. Many of these variants have intermediate sensitivity to drugs as well as intermediate levels of fitness. The development of resistance is thus not all or none, but a continuous evolution of the bacterial membrane, which is qualitatively different from the resistance when targeting enzymes.

### 4.3 Fitness can not be regained without loss of resistance

When under the pressure of a conventional drug, an enzyme mutates, the fitness can be regained by a compensatory mutation while still maintaining the drug resistance.^54^ But in case of AMPs, the factor that determines the resistance or the fitness is the membrane phospholipid composition. In the presence of AMPs, SNP in MprF results in an increase in %LPG, inadvertently compromising the fitness. To maintain homeostasis, the bacterium has to reverse the charge modifications which will also reverse the resistance, resulting in a higher death rate. When using AMPs, the fitness is intricately and inversely coupled to the drug resistance. A second mutation in MprF is more likely to cause a loss of lysylation that helps regain fitness. From our model, it does not seem easy to compensate for the loss of fitness and to recover the viability.

### 4.4 Reversal differentiates qualitatively

While resistance is not instantaneous, the loss of MprF factor upon drug withdrawal is almost instantaneous,^55^ getting affected within a few cell divisions. This *ad hoc* nature of the drug resistance, where the resistance appears under AMP drug pressure and relaxes when the drug pressure is relaxed is qualitatively different from the drug resistant that can remain persistent due to compensatory mutations. The intricate competition between fitness which can not be regained by compensatory mutations forces the bacterium to revert to a drug-sensitive stage. Other models of drug resistance such as by the release of peptidases have not been considered in this work, as they are considered secondary to the surface modifications. Other secondary resistance mechanisms and the chance that the resistance to membrane active drugs remains persistent because of other mechanisms need to be studied as a followup.

## Conclusions

In this work, we studied in detail the mechanism of development of resistance to AMPs with lysylation of PG that follows as a consequence of MprF mutations. The trade-offs between the homeostasis and the antibiotic resistance were evaluated. The model clearly shows a two-step development of resistance when targeting their membranes, where a mutation and an acceleration of the lysylation of membrane lipids both have to happen. This two step development of resistance makes it slower to develop antibiotic resistance against AMPs. Further the apparent lack of compensatory effects suggests to retain the sensitivity of the bacteria to the antibiotics when the drug pressure is relaxed. The relative effects of other mechanisms such as the addition of alanine in teichoic acid or by AMP efflux systems need to be evaluated as well in future studies. If indeed MprF mutation is the most relevant mechanism of resistance development against AMPs, it remains to be seen whether having a secondary drug targeting MprF will be an effective solution to reduce the chances of drug resistance.

---

**Supplementary Figure 1:**
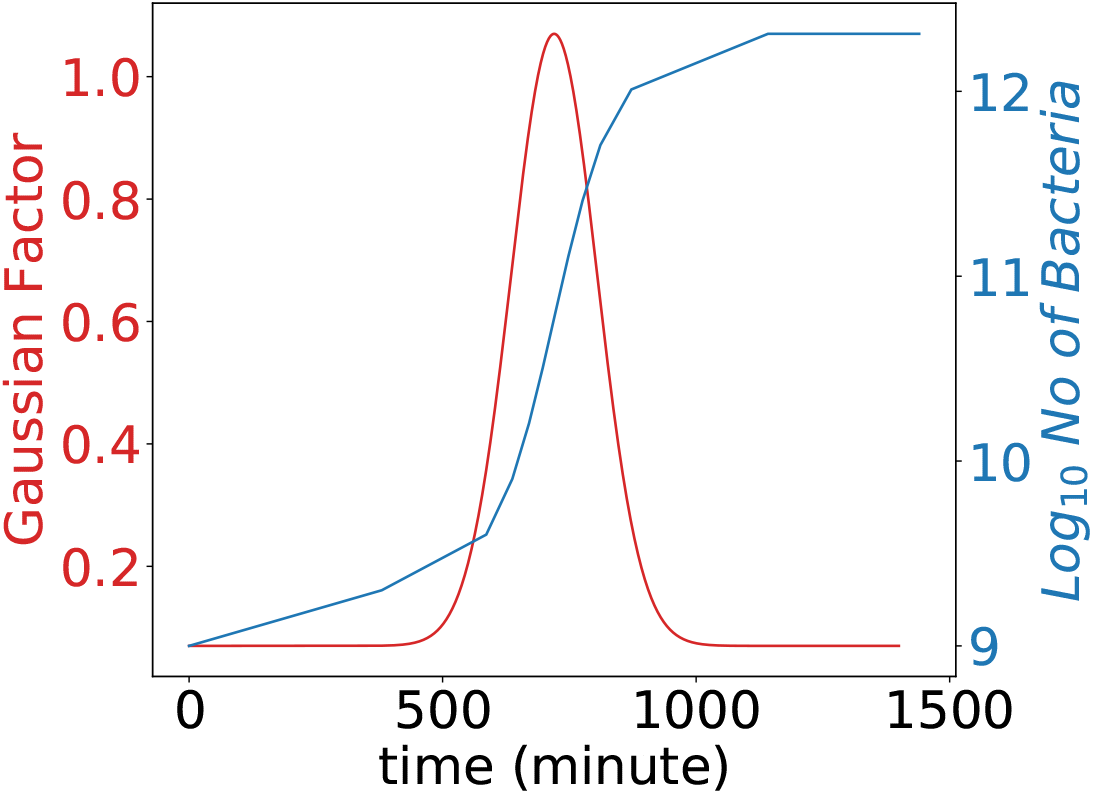
The increase in the number of bacterial cells over time, and the gaussian factor which is supposed to capture the phase dependent variation in growth rate are both shown over one single passage lasting a day. The choice of the gaussian is motivated by the fact that the growth rate is a derivative of the population (sigmoidal) over time.

**Supplementary Figure 2:**
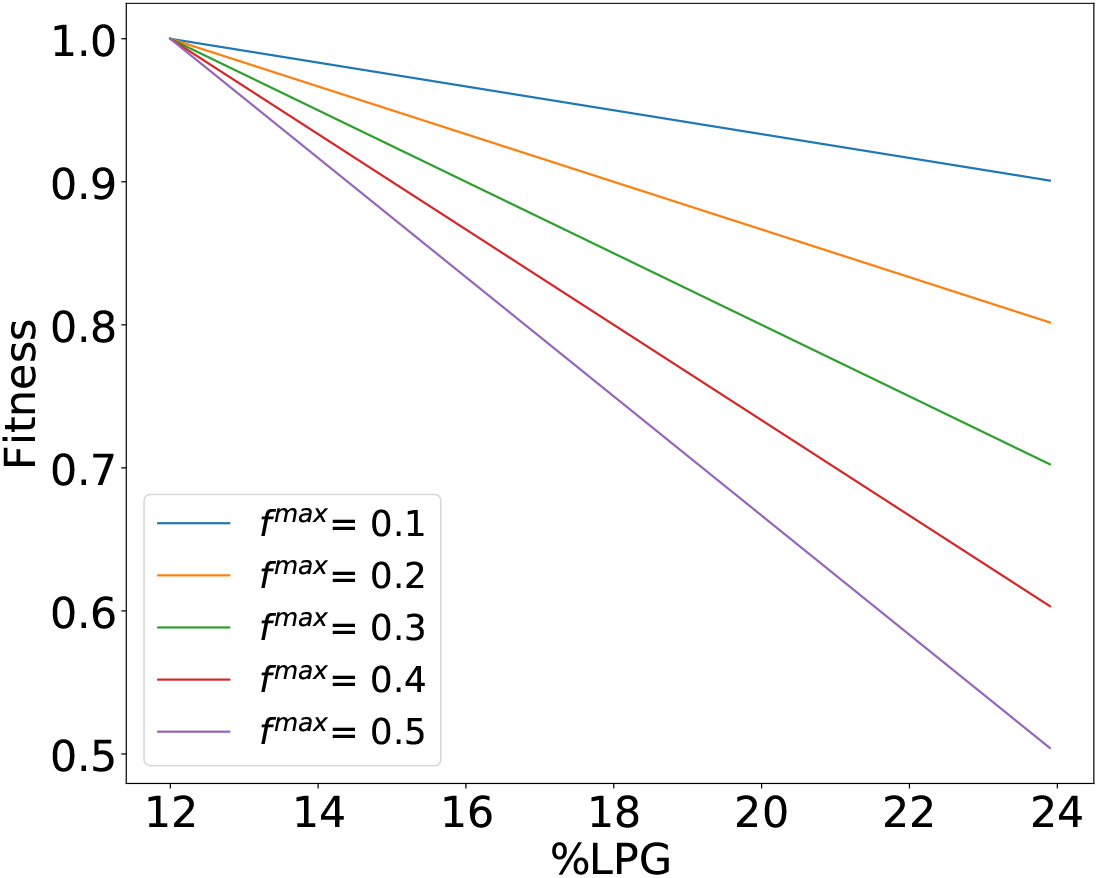
A drop in fitness of the mutant cells which was linear with the change in %LPG was assumed. The parametric dependence on *f^max^* is also shown.

**Supplementary Figure 3:**
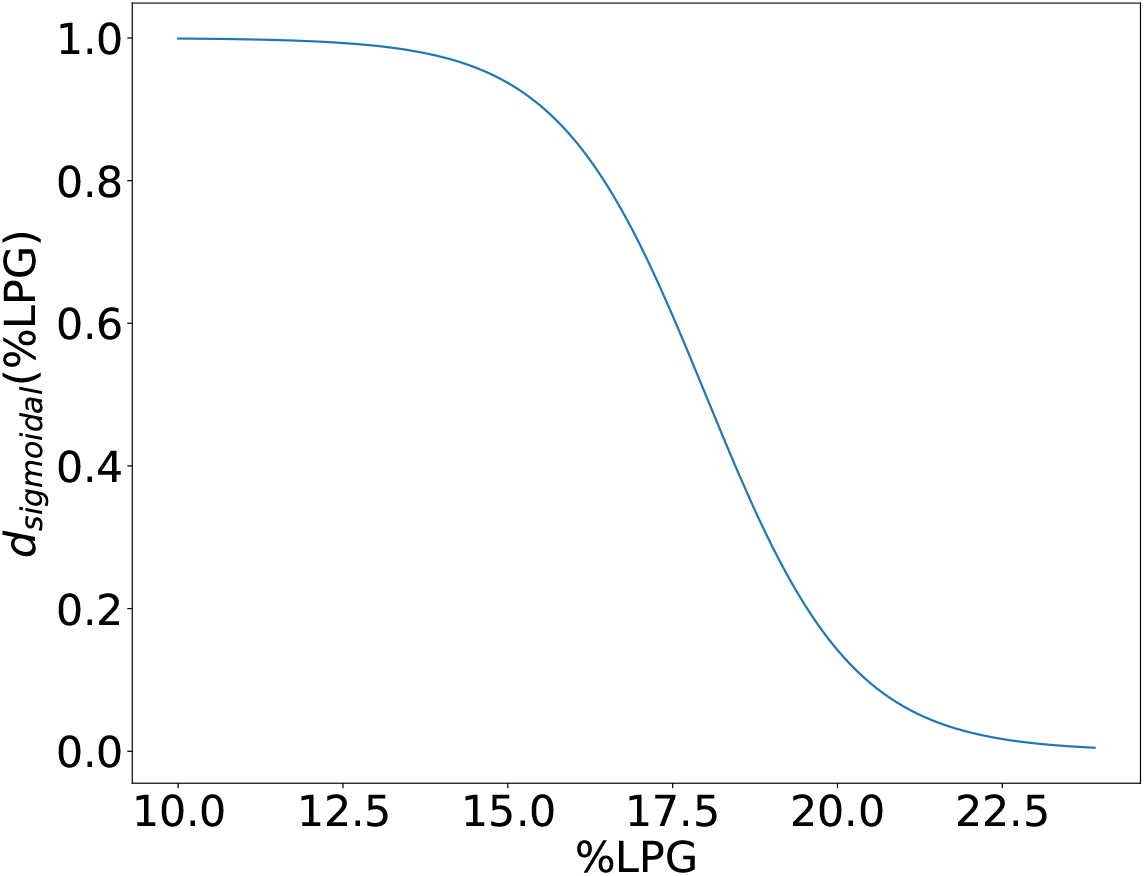
Change in death rate (*d_sigmoidal_*(%*LPG*)) due to the increase in LPG was modeled with a sigmoid, which is shown here.

**Supplementary Figure 4:**
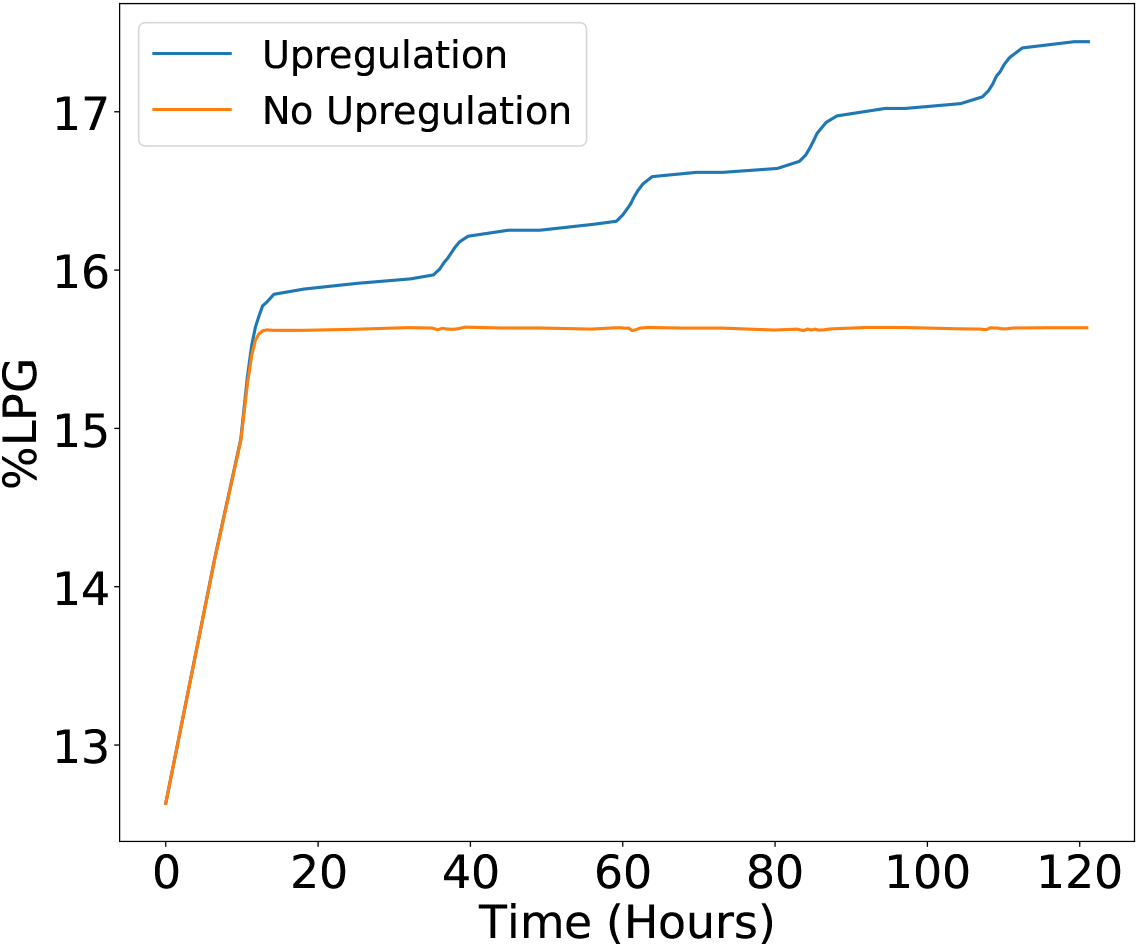
Change of %LPG in mutant cells with time. Results obtained without considering the upregulation of MprF is represented by orange color line and considering the 0.9% upregulation of MprF (*f_MprF_* = 1.009) after each cell division is represented by blue color line.

**Supplementary Figure 5:**
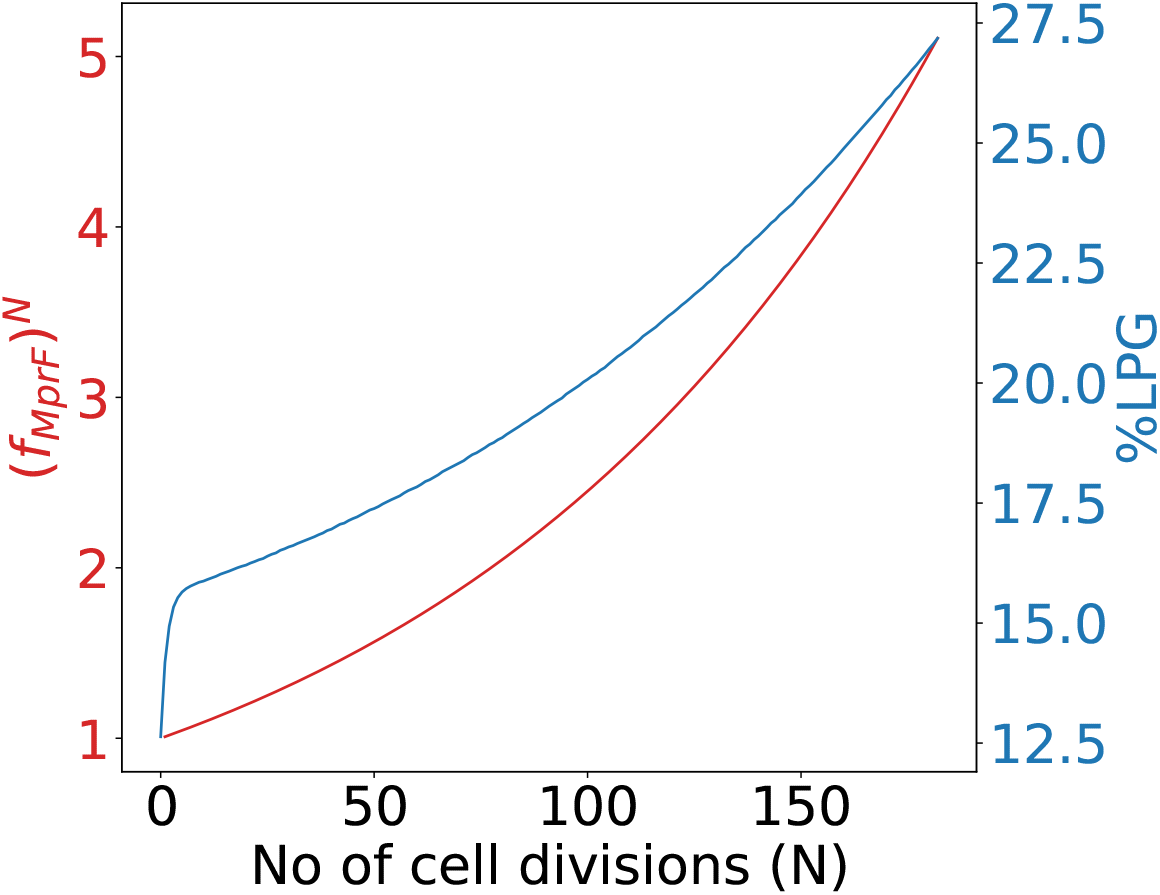
Change in MprF multiplication factor and %LPG of a cell with total number of cell divisions (*N*), by considering an upregulation, *f_MprF_* = 1.009, after every cell division.

**Supplementary Figure 6:**
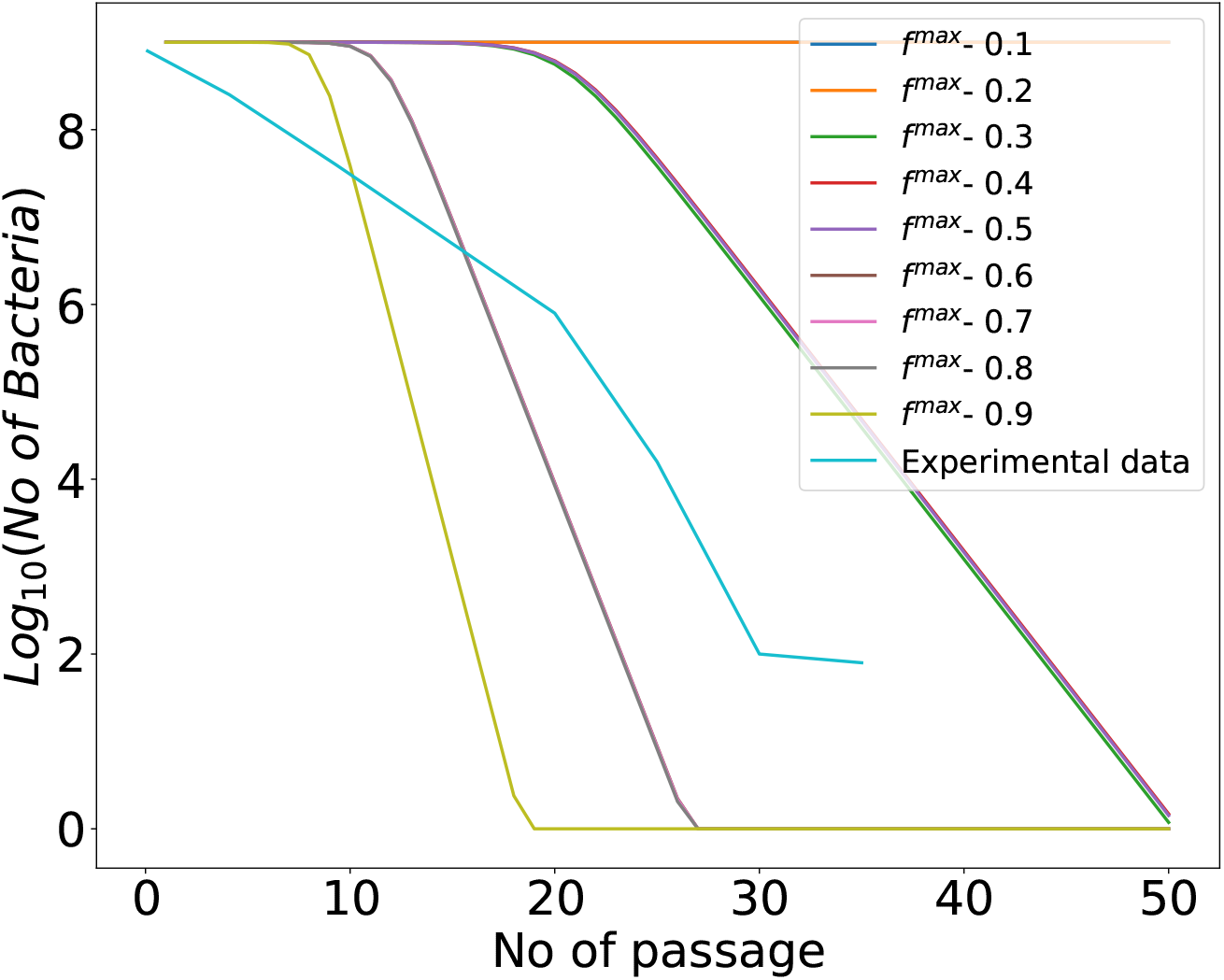
Change in the number of mutant cells over the serial passages in absence of AMP, starting with 10^9^ mutant cell, by considering different values of *f^max^*. Considering the rate at which the bacterial numbers decrease, *i.e*., the slope of the curves, *f^max^* = 0.3 was used for obtaining the results discussed in this work.

